# Addressing viral genomic variability towards developing a Cas13b-based therapy

**DOI:** 10.64898/2026.07.16.739033

**Authors:** Mai Anh Thu Le, Laura C. McCoullough, Zak T. Janetzki, Yi Wen Liaw, Mohamed Fareh, Joseph A. Trapani, Peter A. Revill, Margaret Littlejohn

**Affiliations:** Victorian Infectious Diseases Reference Laboratory, Royal Melbourne Hospital, at the Peter Doherty Institute for Infection and Immunity, Melbourne, Victoria, Australia; Department of Infectious Diseases, The University of Melbourne at the Peter Doherty Institute for Infection and Immunity, Melbourne, Victoria, Australia; Cancer Immunology Program, Peter MacCallum Cancer Centre, Melbourne, Victoria, Australia; Sir Peter MacCallum Department of Oncology, The University of Melbourne, Parkville, Australia

## Abstract

Viral genome diversity may limit the effectiveness of antiviral RNA-editing tools such as CRISPR-Cas13 that can be used to destroy specific mRNA targets, by introducing mismatches between viral RNA targets and CRISPR guide RNAs (crRNAs). These mismatches can reduce target recognition and cleavage efficiency, diminishing antiviral activity and increasing the risk of viral escape. The extent to which natural viral genomic variability limits CRISPR-Cas13 efficacy remains unclear. Here, we used hepatitis B virus (HBV), which has substantial genetic diversity, as a model to assess the impact of viral genome variation on Cas13b activity *in vitro*. The efficacy of *Psp*Cas13b was examined across six HBV genotypes and sub-genotypes using five crRNAs that had up to five mismatches to the target region. We showed that crRNAs with one mismatch to the target strongly suppressed viral antigen expression for all genotypes tested, while some crRNAs with three or more mismatches were less effective. Restoring complementarity using spacer-target mutagenesis improved the level of knockdown for some but not all HBV genotypes, suggesting that sequence specificity alone did not control *Psp*Cas13b efficacy. Our findings show that a “one size fits all” approach for *Psp*Cas13b-mediated treatment of HBV is unlikely to be effective, but the impact of sequence variability on *Psp*Cas13b efficacy can be readily addressed through appropriate design of crRNAs. This approach will likely be necessary for all viral pathogens with highly variant genomes.

**IMPORTANCE:** CRISPR-Cas13 is being explored as a novel antiviral for several viral infections. Viral sequence divergence can compromise CRISPR-Cas13 efficacy by introducing mismatches between therapeutic guide RNAs and viral targets. However, the impact of naturally occurring viral genomic variation on CRISPR-Cas13 efficacy remains poorly understood. Using hepatitis B virus (HBV) as a model, we showed that the effect of mismatches on Cas13b efficacy was context-dependent and varied for different crRNAs, HBV genotypes and target sites. Restoring complementarity improved the efficacy for some, but not all crRNAs, suggesting that Cas13b efficacy was not solely influenced by the number of mismatches. As the target sequence may differ between viral variants, this study advances our understanding of the impact of mismatches on Cas13b efficacy and provides further insights into using Cas13b as a novel antiviral.

## INTRODUCTION

Recent advances in gene-editing, particularly Clustered Regularly Interspaced Short Palindromic Repeats (CRISPR-) Cas technologies, have demonstrated their strong antiviral potential due to the ability to directly target viral gene expression (1–6). CRISPR-Cas13 is a Class II Type VI effector protein that specifically targets single-stranded RNA (ssRNA) (7). CRISPR-Cas13 activation relies on a CRISPR guide RNA (crRNA) comprising a direct repeat sequence and a spacer sequence (8). The direct repeat consists of a stem-loop structure that is essential for the binding of the crRNA to the Cas13 nuclease, forming a ribonucleoprotein complex (9). The spacer sequence is a programmable region, typically 22-30 nucleotides in length, depending on the Cas13 orthologue (9, 10). Activation of Cas13 nuclease requires complementary hybridization between crRNA spacer and a target ssRNA, enabling precise target recognition and specific cleavage of the target ssRNA at a post-transcriptional level. Due to its high programmability and specificity in genome editing, CRISPR-Cas13 is a versatile platform that could be used to treat multiple viral infections (1–6).

Sequence mismatches are known to be important barriers to Cas13 activation (6, 11–17). Previous *in vitro* studies have shown that for *Prevotella species (Psp)* Cas13b, more than four random non-consecutive mismatches between the crRNA and target site completely impairs *Psp*Cas13b activity, by disrupting target recognition and impairing target loading into the ribonucleoprotein complex nuclease (11). In addition, *Psp*Cas13b can tolerate a single nucleotide mismatch that arises from viral variants (6). However, the impact on *Psp*Cas13b activity of more than one naturally occurring mismatch that arises due to viral genetic diversity is unclear. Hepatitis B virus (HBV) has extensive genetic diversity which we have used as a model to test the impact of naturally occurring mismatches in the target site on *Psp*Cas13b efficacy. HBV is classified into ten genotypes (A-J), several of which are further divided into sub-genotypes, distinguished by 8% and 4% nucleotide (nt) divergence respectively, with distinct geographic distributions, natural history and treatment response (18, 19). In a recent proof-of-concept study, *Psp*Cas13b effectively suppressed HBV antigen and virion production in pre-clinical models (5). Multiple crRNAs, binding to different highly conserved regions of HBV RNA transcripts (**Figure 1A**), reduced HBV antigen expression using *in vit*ro and *in vivo* models by 90% and 50% respectively. In this study, both *Psp*Cas13b crRNAs 5 and 9 demonstrated poly-genotypic effects, efficiently reducing antigen secretion for HBV genotypes A2, B2, C2, D3 and E (5). The poly-genotypic effect of additional crRNAs was not assessed. Notably, all crRNAs in this study were designed to be fully complementary to the HBV genotype D3 reference genome. Among the crRNAs tested, both crRNA 5 and 9 had up to a single base mismatch to the genotypes tested, but alignment of the corresponding crRNA target sites across patient samples revealed more extensive sequence variations, with multiple mismatches at various positions (5). The impact of the natural sequence variation inherent in multiple viral genotypes on the antiviral activity of *Psp*Cas13b in the setting of HBV infection is unclear. This is important as HBV infection remains a major global health challenge, with an estimated 254 million people living with chronic hepatitis B worldwide (20), which can result in cirrhosis and hepatocellular carcinoma. There is no cure for chronic HBV infection and new direct acting antiviral approaches are urgently required. Although a DNA virus, HBV replication is mediated via an RNA intermediate that, along with viral mRNAs, is an attractive target for RNA-targeted therapy.

**Figure 1:**
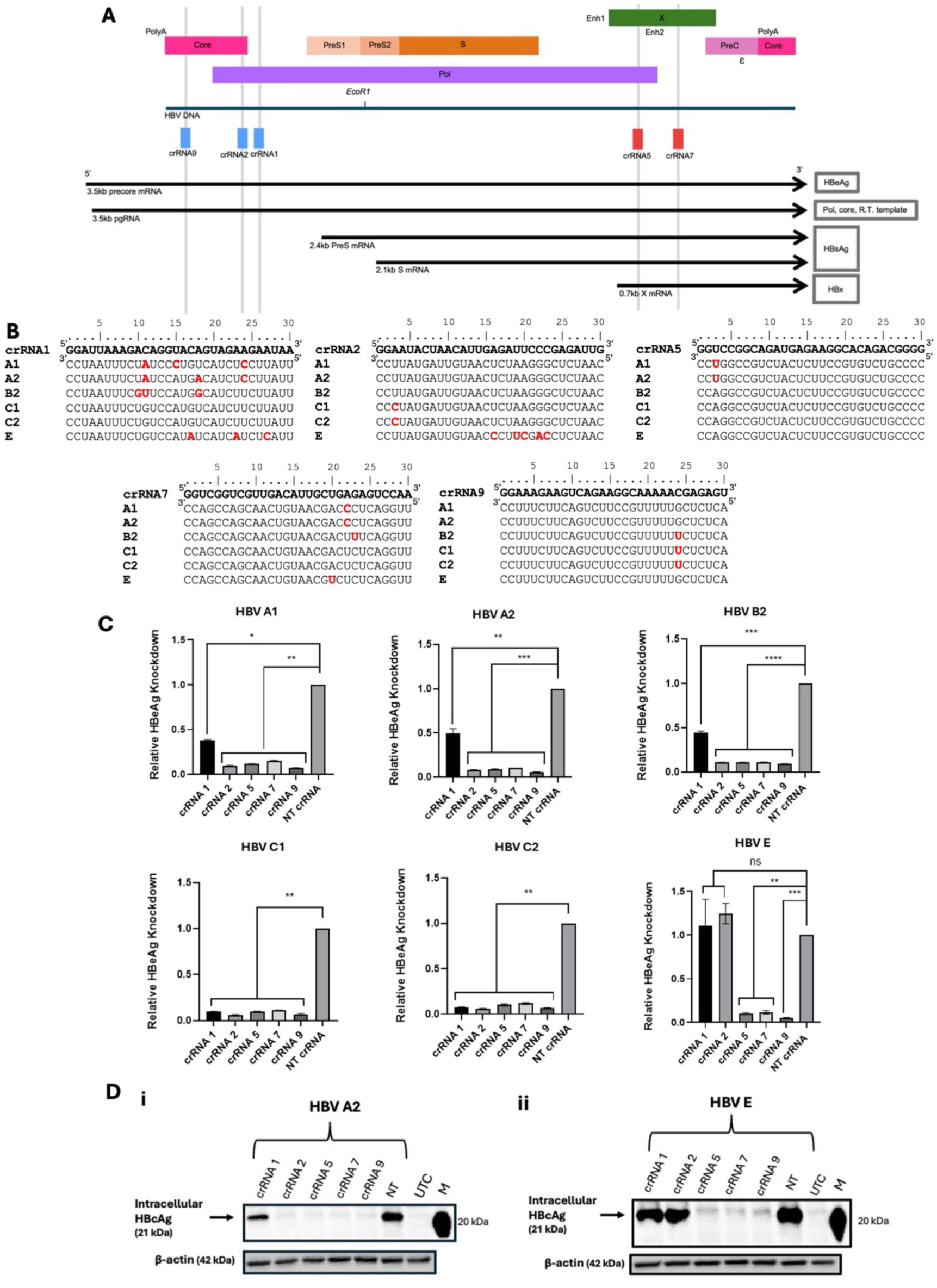
*Psp*Cas13b crRNAs 5, 7 and 9 were effective against all HBV genotypes tested, whereas crRNA 1 and 2 were less effective against different HBV genotypes. **(A)** Map of the target sites of each crRNA 1, 2, 5, 7, and 9 on HBV RNA transcripts. crRNAs 1, 2 and 9 target different conserved regions at the 5’ end of pregenomic RNA (pgRNA) and precore mRNA (pcRNA), which encodes HBcAg and HBeAg respectively. crRNAs 5 and 7 target the regions at the common 3’ end of all HBV RNA transcripts (5). **(B)** Sequence alignments of spacer regions of crRNAs and their respective targets across the six HBV genotype and sub-genotype clones used in this study. Red nucleotide indicates a mismatch. **(C)** Relative secreted HBeAg knockdown compared to the NT crRNA control from HepG2 cells co-transfected with 1.3-mer HBV plasmids encoding genotypes A1, A2, B2, C1, C2 or E, *Psp*Cas13b-BFP plasmid, and crRNA-expressing plasmid. Error bars show SEM and *p*-values following unpaired T-tests on raw data (*P ≤ 0.05, **P ≤ 0.01, *** P ≤0.001, **** P ≤0.0001, ns = p>0.05), n=3, n is the number of independent biological replicates. **(D)** Western blot of intracellular HBcAg protein harvested from transfected HepG2 cells 5 days post-transfection, probed with the in-house 1D8 antibody. crRNAs, Cas13b CRISPR RNAs; HBeAg, HBV e antigen; HBsAg, HBV surface antigen, HBcAg, HBV core antigen; M, marker; NT, non-targeting; UTC, untransfected control.

Building on prior findings (5, 6, 11), this study explored the efficacy of *Psp*Cas13b against diverse HBV genotypes and sub-genotypes, including the underrepresented African genotypes A1 and E. We also analysed additional HBV-targeting crRNAs to evaluate the impact of sequence mismatches between crRNAs and target viral sequences on *Psp*Cas13b efficacy. We showed that crRNAs with up to one mismatch were effective for all HBV genotypes tested, and although some crRNAs with three or more mismatches had reduced efficacy, this was dependent on the crRNA and/or HBV genotype. The nucleotide position of mismatches did not influence *Psp*Cas13b efficacy, suggesting other mechanisms that reduced *Psp*Cas13b activity may be involved. Together, these findings will inform future development of pan-genotypic RNA-editing antiviral strategies.

## RESULTS

### Spacer-target mismatches differentially influenced *Psp*Cas13b activity across HBV genotypes

Since all crRNAs were designed to be fully complementary to HBV genotype D3 (5), all crRNAs had one or more mismatches with at least one of the HBV genotypes included in this study. crRNA 1 had three mismatches at various positions with genotypes A1, A2, B2, and E, while preserving complete spacer-target complementarity with genotypes C1 and C2. crRNA 2 had five mismatches with genotype E, a single mismatch with C1 and C2, and was fully complementary to genotypes A1, A2 and B2. crRNAs 5, 7, and 9 displayed at most a single nucleotide mismatch with all genotypes tested (**Figure 1B**).

#### Impact of sequence mismatches on HBV antigen expression

All *Psp*Cas13b-crRNAs targeted the precore mRNA which expresses the secreted hepatitis B e antigen (HBeAg) (**Figure 1A**), however co-transfection with *Psp*Cas13b-crRNAs differentially impacted HBeAg expression relative to the NT crRNA control across HBV genotypes and sub-genotypes (**Figure 1C**). crRNAs 5, 7, and 9, which had at most one mismatch with the target HBV sequence had greatest knockdown effect, suppressing HBeAg secretion by approximately 88-95% across all genotypes and sub-genotypes tested. In contrast, crRNA 1 which had up to three mismatches with the HBV target, had varied effect on HBeAg expression. For genotypes with complete complementarity (C1 and C2), HBeAg expression was suppressed by 85-90%.

However, crRNA 1 was less effective against sub-genotypes A1, A2, and B2 that had three mismatches, reducing secreted HBeAg by 62%, 50%, and 55% respectively. Interestingly, crRNA 1 had no effect on secreted HBeAg expression for HBV genotype E, despite also encoding three mismatches. The locations of the three mismatches varied for each genotype, although two of the mismatches in the target site of sub-genotype A2 were located in identical positions for sub-genotypes A1 and B2. The target site for crRNA 2 was highly conserved across HBV genotypes and suppressed HBeAg expression for all genotypes except genotype E, with which it had five mismatches.

The impact of crRNAs on the expression of intracellular hepatitis B core antigen (HBcAg), which forms the viral nucleocapsid and is critical for viral replication, was also assessed for genotypes A2 and E. crRNAs 5, 7 and 9 (at most one mismatch) strongly suppressed HBcAg expression in both genotypes. crRNA 2 only suppressed HBcAg expression in genotype A2 (complete complementarity) but had no impact on HBcAg expression for genotype E (five mismatches) (**Figure 1D**). crRNA 1 was less effective in inhibiting HBcAg expression for both genotypes A2 and E (three mismatches), aligning with the secreted HBeAg analysis and further confirming the impaired activity of crRNAs 1 and 2 for these genotypes.

### Mismatches impaired crRNA 1, but not crRNA 5,-mediated knockdown of HBV antigen expression

To determine whether the three spacer-target mismatches for sub-genotypes A1, A2, B2 and E were responsible for the variable efficacy of *Psp*Cas13b crRNA 1, complementarity between the crRNA and target site was restored (**Figure 2A**). This generated four different crRNA 1 variants, designated G1ʹA1, G1ʹA2, G1ʹB2, and G1ʹE, which were 100% complementary to HBV sub-genotypes A1, A2, B2, and E, respectively (**Figure 2B, Table S2**).

**Figure 2:**
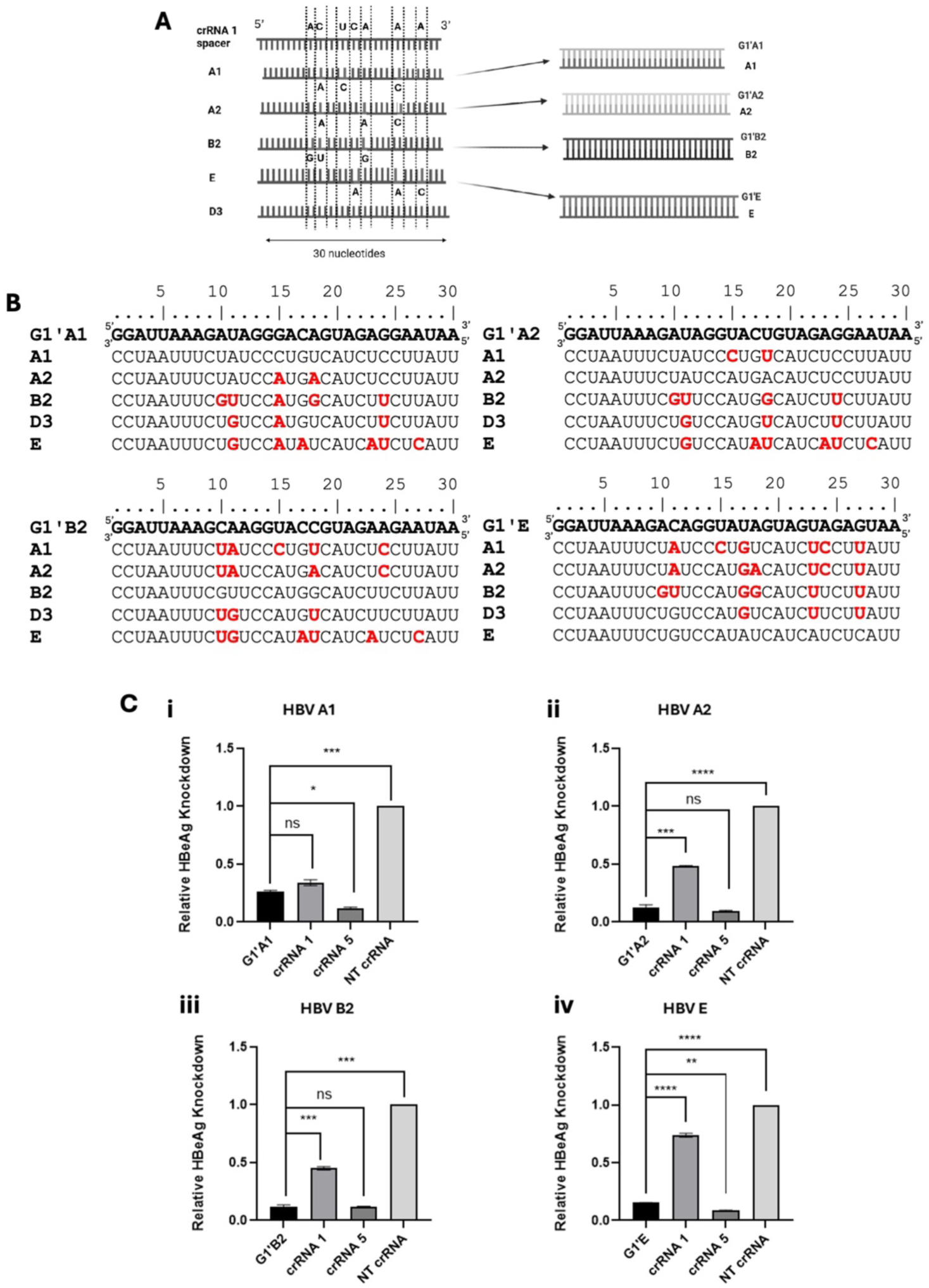
Restoration of crRNA 1 complementarity to different HBV genotypes improved efficacy for some genotypes. **(A)** Schematic of the generation of modified crRNA 1 guides to restore complementarity for HBV genotypes A1, A2, B2 and E. **(B)** Sequence alignments of spacer regions of modified crRNA1 guides and their respective targets. Red nucleotide indicates a mismatch. **(C)** Relative secreted HBeAg knockdown of G1’ crRNAs, crRNA 1 and crRNA 5 compared to the NT crRNA across multiple HBV genotypes from HepG2 cells co-transfected with 1.3mer HBV plasmids of genotypes A1, A2, B2, or E (Panels i-iv), *Psp*Cas13b-BFP plasmid, crRNA-expressing plasmid encoding G1’A1/A2/B2/E crRNAs, crRNA 1, crRNA 5 or NT crRNA. Error bars show SEM and *p*-values following unpaired T-tests on raw data (*P ≤ 0.05, **P ≤ 0.01, *** P ≤0.001, **** P ≤0.0001, ns = p>0.05), n=3, n is the number of independent biological replicates. crRNAs, Cas13b CRISPR RNAs; HBeAg, HBV e antigen; NT, non-targeting.

The knockdown efficacy of the modified G1ʹ crRNAs for their respective fully matched genotypes was compared to the unmodified crRNA 1. The modified G1ʹA2, G1ʹB2, and G1ʹE crRNAs increased knockdown activity compared to the unmodified crRNA 1 for genotypes A2, B2 and E, but not sub-genotype A1. Elimination of spacer-target mismatches increased the knockdown efficacy of G1ʹA2 and G1ʹB2 crRNAs by approximately 33% relative to the unmodified crRNA 1, with a total HBeAg knockdown of 88% (**Figure 2C**). Restoring spacer-target complementarity improved G1ʹE crRNA HBeAg knockdown for genotype E by 59%, compared to the unmodified crRNA 1. Notably, these G1’ crRNAs achieved comparable efficacy to the potent crRNA 5. However, restoring complementarity for the modified G1’A1 crRNA did not improve *Psp*Cas13b activity for genotype A1, as it suppressed secreted HBeAg expression to a similar degree as the unmodified crRNA 1.

To understand how mismatches at different target regions influence *Psp*Cas13b efficacy, the crRNA 5 target site was further investigated, due to its demonstrated strong poly-genotypic efficacy (5). crRNA 5 targets all HBV RNAs (**Figure 1A**). crRNA 5 had an inherent single base mismatch with sub-genotypes A1 and A2 and was 100% complementary to sub-genotypes B2 and genotype E (**Figure 1B**). Three additional mismatches were introduced into crRNA 5 to generate four modified G5’ variants: G5ʹA1, G5ʹA2, G5ʹB2, and G5ʹE (**Figure 3A**). The newly introduced mismatches for each G5’ crRNA were based on the positions of the crRNA 1 mismatches for individual genotypes A1, A2, B2, and E (**Figure 3B** & **Table S3**). The efficacy of each modified G5’ crRNA, encoding three to four mismatches at various positions, was compared to the unmodified crRNA 5 and the NT control for the corresponding genotype (A1, A2, B2, E, or D3), by measuring the level of secreted HBeAg, hepatitis B surface antigen (HBsAg) and intracellular HBcAg expression in transfected cells.

**Figure 3:**
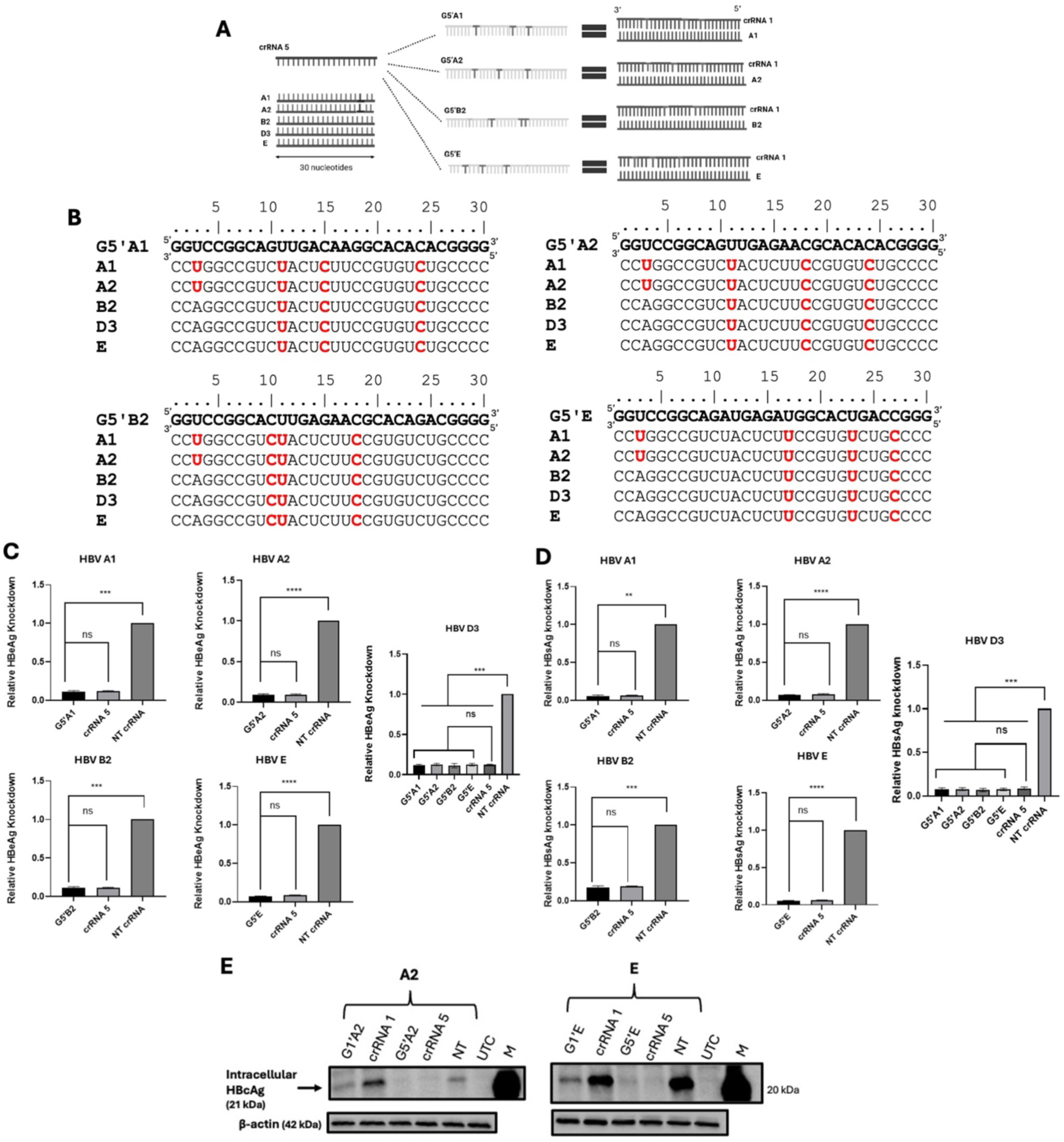
The introduction of three to four mismatches did not impact crRNA 5 efficacy. **(A)** Schematic of the generation of modified crRNA 5 guides to introduce mismatches at the same positions as the crRNA 1 mismatches for HBV genotypes A1, A2, B2 and E. **(B)** Sequence alignments of spacer regions of modified crRNA 5 guides and their respective targets. Red nucleotide indicates a mismatch. Relative secreted **(C)** HBeAg and **(D)** HBsAg knockdown compared to the NT crRNA control from HepG2 cells co-transfected with plasmids expressing 1.3mer HBV of genotype A1, A2, B2, E or D3, *Psp*Cas13b-BFP, and crRNAs G5’A1, G5’B2, G5’E, crRNA 5 or NT control. Error bars show SEM and *p*-values following unpaired T-tests on raw data (*P ≤ 0.05, **P ≤ 0.01, *** P ≤0.001, **** P ≤0.0001, ns = p>0.05). n=3, n is the number of independent biological replicates. **(E)** Western blotting of intracellular HBcAg protein harvested from transfected HepG2 cells 5 days post-transfection, probed with the in-house 1D8 antibody. crRNAs, Cas13b CRISPR RNAs; HBcAg, HBV core antigen; HBV e antigen; M, marker; NT, non-targeting, UTC untransfected control.

Despite having three to four mismatches, all G5ʹ variants reduced HBeAg to similar levels as the unmodified crRNA 5 for all HBV genotypes tested (A1, A2, B2, and E) (**Figure 3C**). In contrast to crRNA 1, where similar mismatch patterns substantially reduced target knockdown for these specific genotypes (**Figure 1C**), the modified G5ʹ crRNAs were not impacted by three or four random mismatches, retaining a strong poly-genotypic effect. Furthermore, when tested against genotype D3, positional variations in the mismatches among the G5ʹ crRNAs did not impact their HBeAg knockdown efficacy (**Figure 3C**).

In a similar manner, all G5’ crRNAs induced strong inhibition of HBsAg secretion across HBV genotypes (**Figure 3D**), underscoring the high tolerance to three to four mismatches of this target site.

The knockdown of intracellular HBcAg by G1’ crRNAs, G5’ crRNAs, and unmodified crRNA 1 and crRNA 5 for HBV genotypes A2 and E, was also compared (**Figure 3E**). Consistent with the previous analysis (**Figure 1D**), the unmodified crRNA 1 encoding three mismatches had reduced efficacy in inhibiting HBcAg expression in genotypes A2 and E, compared to the fully complementary, modified G1’A2 and G1’E crRNAs (**Figure 3E**). On the contrary, G5’A2 and G5’E crRNAs, each encoding three mismatches at the same positions as crRNA 1 for each respective genotype, achieved strong intracellular HBcAg knockdown, with suppression levels comparable to the unmodified, fully complementary crRNA 5, for both A2 and E genotypes (**Figure 3E**).

### Differences in mismatch positions alone did not account for the variable knockdown efficacy of crRNA 1 across genotypes

Modifying crRNA 1, which was fully complementary to genotype D3 (5), to generate the G1ʹ variants (A1, A2, B2, E) introduced concomitant mismatches to the HBV D3 target (**Table S4**, **Figure 4A**), enabling investigation of the effect of different mismatch positions on *Psp*Cas13b-crRNA activity, rather than just total mismatch number. The relative HBeAg knockdown of the modified G1’ crRNAs against sub-genotype D3 were compared.

**Figure 4:**
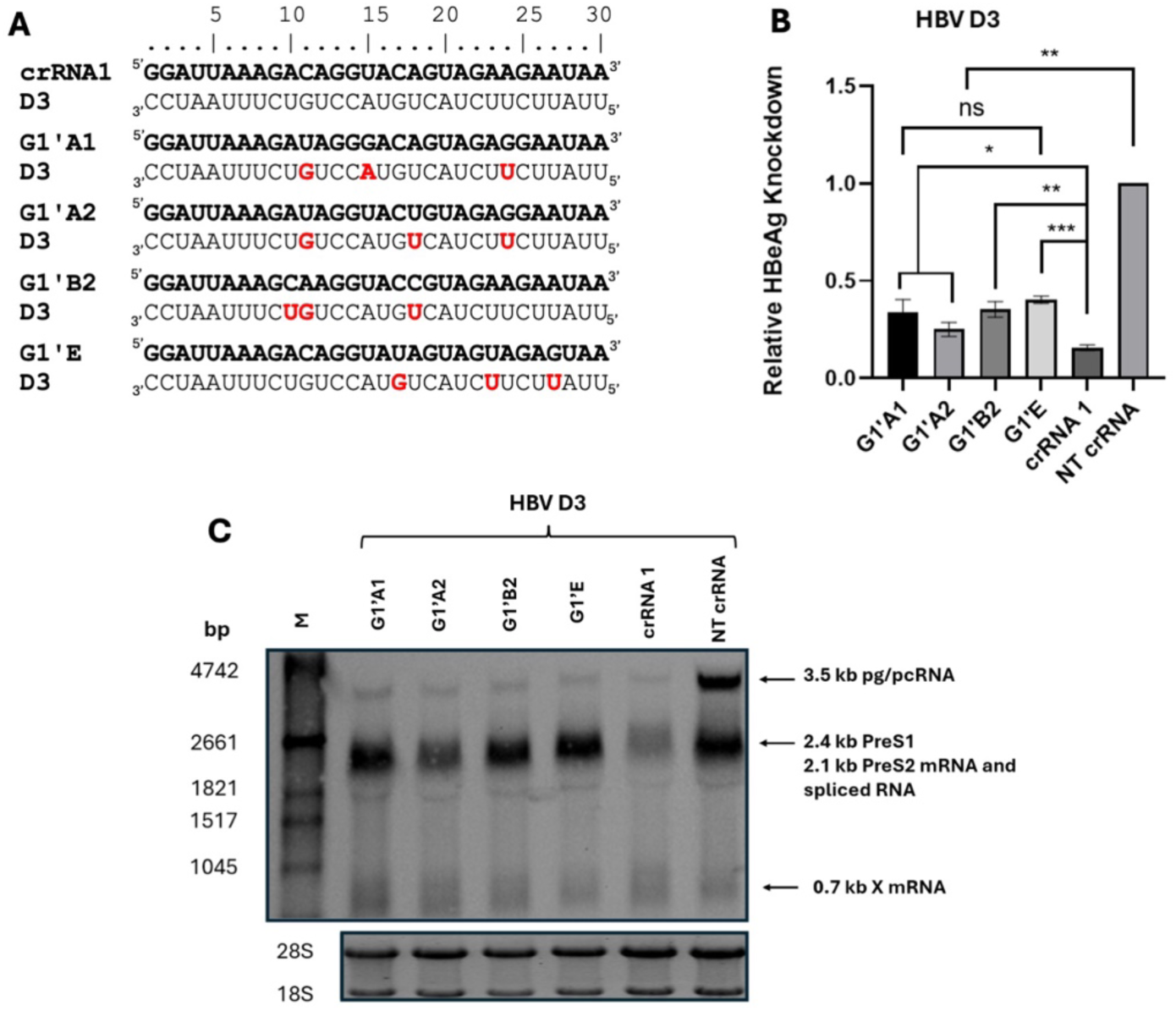
Mismatch position did not influence *Psp*Cas13b activity. **(A)** Sequence alignments of spacer regions of modified crRNA1 guides against HBV genotype D3. Red nucleotide indicates a mismatch. **(B)** Relative secreted HBeAg knockdown compared to the NT crRNA control from HepG2 cells co-transfected with 1.3mer HBV D3 genotype plasmid, *Psp*Cas13b-BFP plasmid, crRNA-expressing plasmid of either crRNAs G1’A1, G1’A2, G1’B2, G1’E, unmodified crRNA 1 or NT control. Error bars show SEM and *p*-values following unpaired T-tests on raw data (*P ≤ 0.05, **P ≤ 0.01, *** P ≤0.001, **** P ≤0.0001, ns = p>0.05), n=3, n is the number of independent biological replicates. **(C)** Northern blotting of HBV RNAs harvested from transfected HepG2 cells 5 days post-transfection. The NT crRNA serves as a negative control, and unmodified crRNA 1 serves as a positive control. HBeAg, HBV e antigen; HBx RNA, HBV X RNA; pg/pcRNA; bp, base pair; crRNA, CRISPR RNA; pg/pcRNA, HBV pregenomic/precore RNA; PreS, HBV pre-surface; NT, non-targeting, M, marker.

Although the modified G1ʹ crRNAs demonstrated significant HBeAg knockdown for sub-genotype D3 relative to the NT control (**Figure 4A**), the modified crRNAs were significantly less effective than the unmodified crRNA 1 that was 100% complementary to sub-genotype D3.

Whilst the unmodified crRNA 1 reduced HBeAg secretion by 85%, G1’A1, G1’A2, and G1’B2 crRNAs, which had mismatches clustered in the centre of the spacer (**Figure 4A**), only suppressed secreted HBeAg by 65%, 75%, and 65% respectively (**Figure 4B**). G1’E crRNA, with mismatches toward the 3’ end of the spacer sequence (**Figure 4A**), also had reduced efficacy compared to the unmodified crRNA 1, with a 60% decrease in secreted HBeAg. Overall, there was no significant difference between the level of HBeAg knockdown for the different modified G1’ crRNAs against genotype D3, despite variation in their mismatch positions (**Figure 4B**).

These findings were validated by northern blot analysis for HBV genotype D3 RNA expression. Unmodified crRNA 1, which was fully complementary to genotype D3 RNA target, suppressed precore/pregenomic RNA (pc/pgRNA) to the greatest extent relative to the NT crRNA (**Figure 4C**). Although the modified G1’ crRNAs suppressed pc/pgRNA compared to the NT crRNA control, they were less effective compared to the unmodified crRNA 1. In general, the levels of pc/pgRNA knockdown were comparable among the G1’ crRNAs.

## DISCUSSION

Mismatches are known to be key barriers to Cas13 target recognition and cleavage, although their impacts can vary remarkably depending on the total number of mismatches within the crRNA-target duplex (6, 11–17). Previous studies that had shown the impact of mismatches on *Psp*Cas13b activity mainly introduced random mismatches into the crRNAs and showed that *Psp*Cas13b can tolerate four or less mismatches without impacting activity (6, 11). Here, we evaluated the impact of naturally occurring, dispersed crRNA-target mismatches arising from natural HBV genomic variability (21) on *Psp*Cas13b activity. We showed that the impact of mismatches was dependent on the crRNA and/or target site, and varied by HBV genotype and sub-genotype. All crRNAs maintained strong activity against genotypes which had a fully complementary target site or encoded one mismatch (**Figure 1**). Consistent with our previous study (5), crRNAs 5 and 9 displayed high efficacy in antigen suppression for multiple genotypes. crRNA 7, which was evaluated for its poly-genotypic activity for the first time, also demonstrated robust target knockdown for all genotypes tested. For crRNA 1, three mismatches partially impaired *Psp*Cas13b activity, and restoration of complementarity improved efficacy for most genotypes tested (**Figure 2**), suggesting that the mismatches caused the reduced efficacy of crRNA 1 for these HBV genotypes, and contrasts with the previous study that showed that *Psp*Cas13b maintains high efficacy against transcripts containing four or less mismatches (11). In contrast, all G5’ variants maintained potent HBV antigen suppression (**Figure 3**), despite encoding three to four mismatches, which aligns with previous findings of *Psp*Cas13b’s resilience to four random mismatches (11). These results indicate that *Psp*Cas13b mismatch sensitivity can be crRNA or target-dependent, with crRNA 5 exhibiting greater intrinsic resilience than crRNA 1. The enhanced mismatch tolerance of crRNA 5 underscores its potential as a strong candidate for Cas13b antiviral development against HBV, capable of achieving a balance between potent activity for diverse viral strains and high specificity.

Restoring complementarity of crRNA 1 did not improve its activity for genotype A1 (**Figure 2**), suggesting that the reduced activity was not solely due to mismatches and may instead be influenced by other underlying viral factors. Sequence divergence between HBV genotypes not only introduces mismatches at the crRNA target site but can also alter local target RNA structure and folding (22), which may potentially hinder target scanning and reduce Cas13b accessibility (11, 23, 24). These genotype-specific structural features may therefore impose additional constraints on Cas13-mediated cleavage, contributing to genotype-specific loss of *Psp*Cas13b efficacy (23, 25, 26). In addition, interaction with viral or host RNA binding proteins at these sites may mask Cas13b recognition. Moreover, the potency of crRNAs can vary significantly with even one-nucleotide shift in the target site (11), suggesting that subtle differences in sequence context or spacer-target environment among the genotypes could further impact *Psp*Cas13b-crRNA cleavage dynamics.

Cas13 sensitivity to mismatches depends on the nucleotide position of the mismatch (6, 11, 17, 27). While the activity of *Pbu*Cas13b, *Lbu*Cas13a, and *Lwa*Cas13a is strongly sensitive to central-spacer mismatches, in positions 12-17, 5-16 and 13-24 respectively (14, 17, 27), the activity of *Psp*Cas13b has minimal sensitivity to central mismatches (positions 14-16) but higher sensitivity to those located near the 5ʹ end (positions 1-2) where the conserved G-G motif resides, determined using artificial reporter systems (6, 11). Our findings using naturally occurring mismatches were consistent with these observations, as we found that variation in mismatch distribution in the central region, specifically mismatches at positions 10, 11, 15, 17, 18, 23, 24 and 27, had minimal impact on the knockdown activity of the *Psp*Cas13b-G1ʹ crRNA against HBV D3 (**Figure 4**). This builds on previous findings that demonstrated that mismatches at positions 7 and 14 had no impact on *Psp*Cas13b activity, whereas a single-nucleotide mismatch at position 21 impaired *Psp*Cas13b activity (11). We also showed that mismatches at identical positions within the same target site produced distinct functional outcomes for *Psp*Cas13b for different HBV genotypes. For instance, three mismatches on crRNA 1 completely abolished *Psp*Cas13b activity against genotype E (**Figure 1C**), whereas mismatches at the identical positions on modified G1ʹE crRNA still resulted in 60% reduction of HBeAg against genotype D3 (**Figure 4B**). These results indicate that spatial distribution of mismatches was not the only driver of variability in the crRNAs’ activity across HBV genotypes, indicating that additional factors may also influence *Psp*Cas13b sensitivity to mismatches. Differences in the nucleotide identities of mismatches can differentially affect crRNA-target duplex stability, potentially contributing to the observed variability in mismatch effects on the system’s activity across HBV genotypes (28, 29). Mismatches between G1ʹE crRNA and the genotype D3 target (G-U, U-U, U-G) primarily include the thermodynamically stable G-U ‘wobble’ pair, which can maintain partial RNA-RNA binding, thereby allowing moderate Cas13b cleavage efficiency (28–30). G-U mismatch was also more tolerated by Cas13 than other single-base mismatches (26). In contrast, the C-A and A-A mismatches observed between unmodified crRNA 1 and the genotype E target are more destabilizing, potentially weakening duplex formation and reducing Cas13b activity through impaired target recognition and loading (28, 31). Thus, the chemical nature of crRNA-target mismatches could be a critical determinant of *Psp*Cas13b mismatch sensitivity and knockdown activity.

Several limitations of this study are acknowledged. Our experiments were performed in HepG2 cells using plasmid transfection, which, although enables high viral replication *in vitro*, does not fully capture the complexity of natural HBV infection or the liver microenvironment. Mismatch profiling was limited to selected crRNAs and patterns, primarily examining mismatch number and position, whilst mismatch identity, target accessibility and potential collateral cleavage were not directly assessed. Despite these limitations, this study provides key insights into factors influencing *Psp*Cas13b activity across diverse HBV strains. By evaluating naturally occurring mismatches between crRNAs and viral targets utilizing the full HBV genome, our work extends beyond the artificial reporter systems employed in previous studies (6, 11–17). Our findings highlight that the number of mismatches that impact *Psp*Cas13b activity is not universal, and may depend on the crRNA sequence, target site and/or mismatch identity.

Overall, this study demonstrates that while mismatches represent a critical challenge for Cas13b-based antiviral therapeutics, their impact is highly context-dependent and can be resolved through careful crRNA design that is personalized to a patient’s dominant viral sequence or multiplexing crRNAs to target different viral sequences. Using lipid nanoparticles and mRNA to deliver Cas13b will permit these strategies. Whilst crRNA 1 demonstrated genotype dependent sensitivity to three mismatches, crRNA 5 exhibited remarkable resilience, retaining strong activity even in the presence of three to four random mismatches in the target site across genotypes, identifying it as a promising candidate for pan-genotypic HBV therapy. Collectively, these findings advance our mechanistic understanding of Cas13b activity and mismatch tolerance in the HBV genome, while also highlighting practical strategies to minimize viral escape and inform the development of effective Cas13b-based therapeutics for viral infections.

## MATERIALS AND METHODS

### 1.3mer WT HBV DNA clones

Greater than genome length 1.3mer WT HBV genotype A1, A2, B2, C1, C2, D3 and E plasmids used in this study have previously been described (32, 33).

### *Psp*Cas13b-NES-HIV-3xFlag-T2A-BFP clone

*Psp*Cas13b-BFP was designed and cloned as previously described (6).

### Design and cloning of *Psp*Cas13b CRISPR RNAs (crRNAs)

*Psp*Cas13b crRNAs 1, 2, 5, 7, 9, and a non-targeting crRNA (NT crRNA) have been previously described (5). The NT crRNA did not have complementarity to any known human or HBV RNA sequence and served as a negative control.

To determine the impact of sequence mismatches on Cas13b crRNA efficacy, the spacer region of crRNA 1 was modified to generate G1’A1, G1’A2, G1’B2 and G1’E to restore complementarity of crRNA 1 to genotypes A1, A2, B2 and E respectively. To determine whether the mismatch positions influenced efficacy, crRNA 5 was modified to generate G5’A1, G5’A2, G5’B2, and G5’E to introduce mismatches in the same positions as the modified crRNA 1 sequences for genotypes A1, A2, B2 and E respectively.

All crRNAs designed in this study were cloned into the pC0043-*Psp*Cas13b crRNA backbone (Addgene #103854, a kind gift from the Feng Zheng lab), using standard molecular cloning techniques as previously described (6).

A map of *Psp*Cas13b crRNAs is provided in Figure 1A and all crRNAs’ spacer sequences are detailed in Table S1.

### Cell culture and transient transfection

HepG2 cells were seeded at 60-80% confluency into 60 mm dishes or 12-well plates. All *in vitro* transfection experiments utilized a combination of three plasmids: the *Psp*Cas13b-BFP plasmid; the HBV 1.3mer wild-type (WT) plasmid; and the crRNA expression plasmid. Each DNA plasmid was transfected at the previously optimized 2:1:28 molar ratio (5), using Lipofectamine 3000 (Thermo Fisher Scientific), according to the manufacturer’s instructions. Transfection efficiency was monitored using a blue fluorescent protein (BFP) expression construct and counting BFP-expressing cells using a fluorescent microscope.

All analyses of intracellular HBV RNA, intracellular protein, and secreted viral proteins (HBeAg and HBsAg) were performed 5 days post-transfection, corresponding to the peak of HBV DNA replication and protein expression (32).

### HBV intracellular protein analysis

Intracellular HBV proteins in cell lysates harvested 5 days post-transfection were detected by immunoblotting as previously described (34), using antibodies against precore/core proteins (in-house monoclonal antibody 1D8) (35–37).

### HBV RNA expression analysis

Total RNA was extracted from transfected cells using the RNeasy® Mini Kit (Qiagen) according to the manufacturer’s instructions. HBV RNAs were analyzed by northern hybridization using the Ambion® NorthernMax™-Gly Kit (Thermo Fisher Scientific), following the manufacturer’s instructions and previously described methods (38).

### Quantitative serology

Secreted HBeAg and HBsAg concentrations in cell culture lysates and supernatants were measured by chemiluminescent microparticle immunoassay using the Roche HBeAg and HBsAg assays on a Cobas e801 instrument as previously described (32).

### Data analysis

Statistical analyses were performed using GraphPad Prism™ Version 7 (GraphPad Software). Data presented in bar graphs represents mean values, with error bars indicating standard error of the mean (SEM). An unpaired two-tailed Student’s *t*-test was used to assess statistical significance between two independent groups. A p-value of less than 0.05 (*p* < 0.05) was considered statistically significant.

## Author Contributions

M.A.T.L performed experiments and generated all graphs and figures. M.A.T.L and L.C.M co-wrote the manuscript and designed the experiments. M.A.T.L and Z.T.J cloned crRNA constructs. L.C.M, Z.T.J and Y.W.L performed quantitative serology. M.F and J.A.T provided the *Psp*Cas13b clone, discussed the project and data and provided advice on experimental design. L.C.M, P.A.R and M.L conceived and supervised the study. All authors read, commented, edited and approved the manuscript.

## Competing Interests

P.A.R has received investigator-initiated industry funded grants from Gilead. None of this support is relevant to the work in this manuscript. The remaining authors declare no conflict of interest.

## Funding Sources

This work was funded by mRNA Victoria through a grant to M.L.

