## Supplementary figures and tables for "Addressing viral genomic variability towards developing a Cas13b-based therapy"

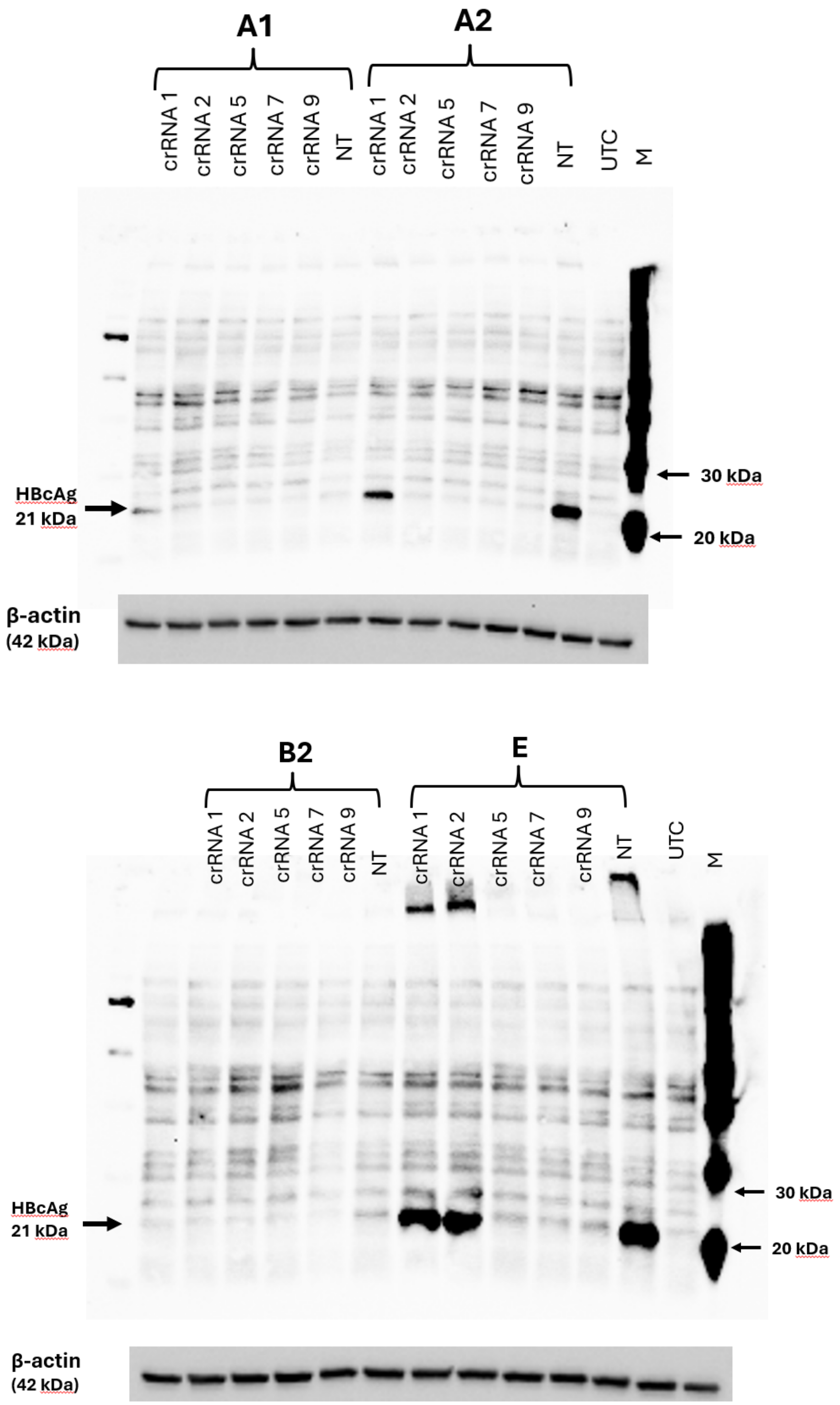


**Figure S1: Full immunoblot showing hepatitis B core antigen (HBcAg) expression in HBV A1, A2, B2 and E genotypes when treated with crRNAs 1, 2, 5, 7 and 9**, from HepG2 cells co-transfected with HBV 1.3 plasmid of genotypes A1, A2, B2, or E, *Psp*Cas13b-BFP, and crRNA-expressing plasmid for crRNAs 1, 2, 5, 7 and 9 or a NT crRNA. BFP, blue fluorescent protein; crRNAs, Cas13b CRISPR RNAs; HBcAg, HBV core antigen; M, marker; NT, non-targeting, UTC untransfected control.

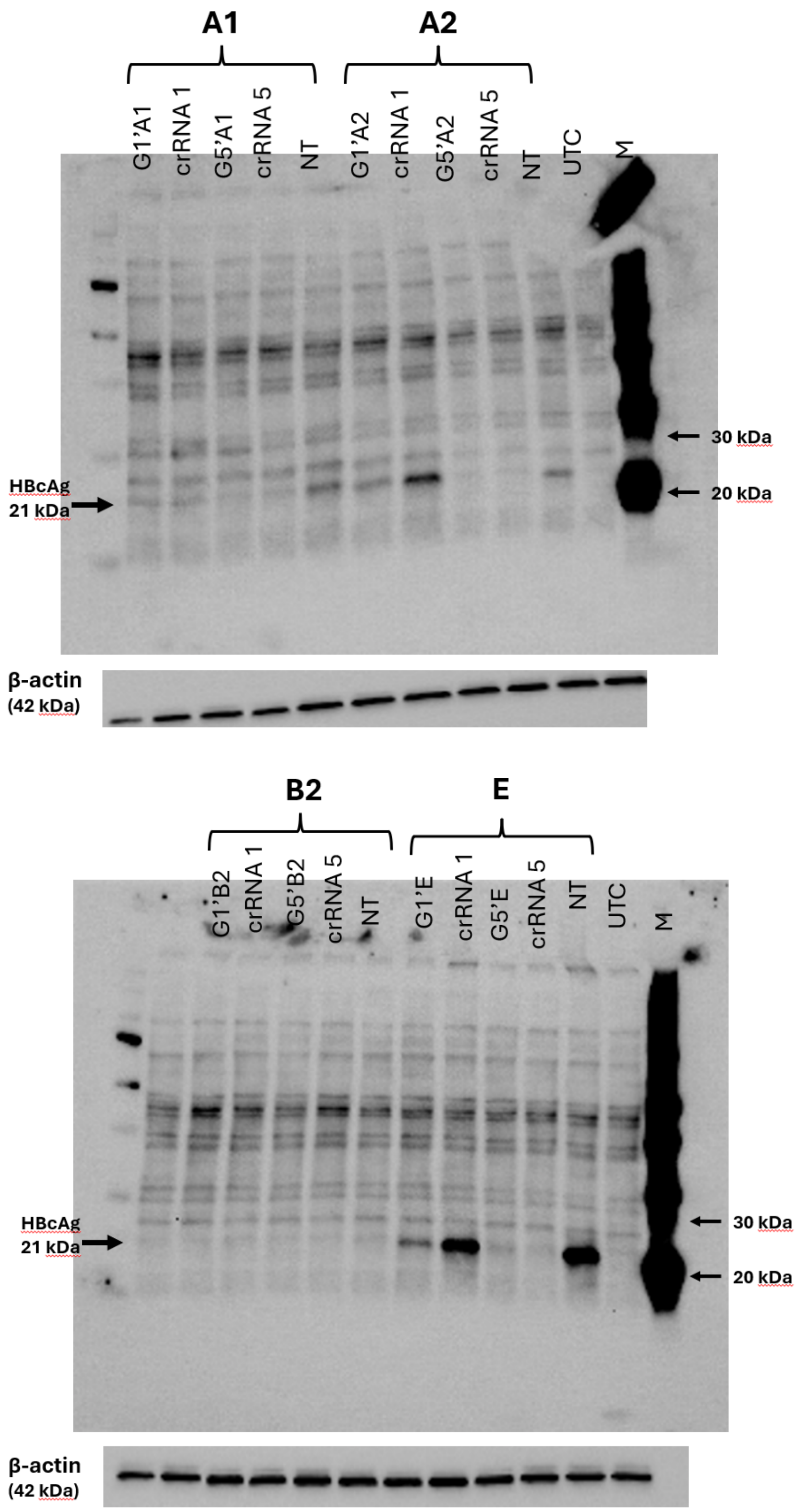


**Figure S2: Full immunoblot showing hepatitis B core antigen (HBcAg) expression in HBV A1, A2, B2 and E genotypes when treated with modified G1' crRNAs, modified G5' crRNAs, unmodified crRNA 1 and unmodified crRNA 5**, from HepG2 cells co-transfected with HBV 1.3

plasmids of genotypes A1, A2, B2, or E, *Psp*Cas13b-BFP, and crRNA-expressing plasmid for modified G1'A1/A2/B2/E crRNAs, modified G5'A1/A2/B2/E crRNAs, unmodified crRNA 1, unmodified crRNA 5 or NT crRNA. BFP, blue fluorescent protein; crRNAs, Cas13b CRISPR RNAs; HBcAg, HBV core antigen; M, marker; NT, non-targeting, UTC untransfected control.

### SUPPLEMENTARY TABLE

**Table S1: Spacer sequences for crRNAs tested.**

| crRNA | Spacer sequence (5'-3') |
|---|---|
| crRNA 1 | GGATTAAAGACAGGTACAGTAGAAGAATAA |
| crRNA 2 | GGAATACTAACATTGAGATTCCCGAGATTG |
| crRNA 5 | GGTCCGGCAGATGAGAAGGCACAGACGGGG |
| crRNA 7 | GGTCGGTCGTTGACATTGCTGAGAGTCCAA |
| crRNA 9 | GGAAAGAAGTCAGAAGGCAAAAACGAGAGT |
| NT crRNA | TAGATTGCTGTTCTACCAAGTAATCCATCA |
| G1'A1 | GGATTAAAGATAGGGACAGTAGAGGAATAA |
| G1'A2 | GGATTAAAGATAGGTACTGTAGAGGAATAA |
| G1'B2 | GGATTAAAGCAAGGTACCGTAGAAGAATAA |
| G1'E | GGATTAAAGACAGGTATAGTAGTAGAGTAA |
| G5'A1 | GGTCCGGCAGTTGACAAGGCACACACGGGG |
| G5'A2 | GGTCCGGCAGTTGAGAACGCACACACGGGG |
| G5'B2 | GGTCCGGCACTTGAGAACGCACAGACGGGG |
| G5'E | GGTCCGGCAGATGAGATGGCACTGACCGGG |

**Table S2: A summary of numbers and positions of mismatches of unmodified crRNA 1 and G1' crRNAs with their targets in HBV genotypes A1, A2, B2 and E**. The positions of mismatches were counted from the 5' end of the crRNA spacer sequence.

| HBV Genotype | crRNAs | Number of mismatches | Positions of mismatches |
|---|---|---|---|
| A1 | crRNA 1 | 3 | 11, 15, 24 |
| | G1'A1 | 0 | |
| | G1'A2 | 2 | 11, 15 |
| | G1'B2 | 5 | 10, 11, 15, 18, 24 |
| | G1'E | 6 | 11, 15, 17, 23, 24, 27 |
| A2 | crRNA 1 | 3 | 11,18, 24 |
| | G1'A1 | 2 | 15, 18 |
| | G1'A2 | 0 | |
| | G1'B2 | 4 | 10, 11,18, 24 |
| | G1'E | 6 | 11, 17, 18 23, 24, 27 |
| B2 | crRNA 1 | 3 | 10, 11, 18 |
| | G1'A1 | 5 | 10, 11, 15, 18, 24 |
| | G1'A2 | 4 | 10, 11, 18, 24 |
| | G1'B2 | 0 | |
| | G1'E | 6 | 10, 11, 17, 18, 23, 27 |
| E | crRNA 1 | 3 | 17, 23, 27 |
| | G1'A1 | 6 | 11, 15, 17, 23, 24, 27 |
| | G1'A2 | 6 | 11, 17, 18, 23, 24, 27 |
| | G1'B2 | 6 | 10, 11, 17, 18, 23, 27 |
| | G1'E | 0 | |

**Table S3: A summary of mismatch positions for the G5' crRNAs with their targets across HBV genotypes A1, A2, B2, D3 and E**. The positions of mismatches were counted from the 5' end of the spacer sequence.

| crRNA | HBV genotype | Positions of Mismatch |
|---|---|---|
| G5'A1 | A1 | 3, 11, 15, 24 |
| | A2 | |
| | B2 | 11, 15, 24 |
| | D3 | |
| | E | |
| G5'A2 | A1 | 3, 11, 18, 24 |
| | A2 | |
| | B2 | 11, 18, 24 |
| | D3 | |
| | E | |
| G5'B2 | A1 | 3, 10, 11, 18 |
| | A2 | |
| | B2 | 10, 11, 18 |
| | D3 | |
| | E | |
| G5'E | A1 | 3, 17, 23, 27 |
| | A2 | |
| | B2 | 17, 23, 27 |
| | D3 | |
| | E | |

**Table S4: A summary of mismatch numbers and positions in unmodified crRNA 1 and modified G1' crRNAs relative to genotype D3 sequence**. The positions of mismatches were counted from the 5' end of crRNA spacer sequence.

| HBV genotype | crRNAs | Numbers of mismatches | Positions of mismatches |
|---|---|---|---|
| D3 | crRNA 1 | 0 | |
| | G1'A1 | 3 | 11, 15, 24 |
| | G1'A2 | 3 | 11, 18, 24 |
| | G1'B2 | 3 | 10, 11, 18 |
| | G1'E | 3 | 17, 23, 27 |
